# YHP: <u>Y</u>-chromosome <u>H</u>aplogroup <u>P</u>redictor for predicting male lineages based on Y-STRs

**DOI:** 10.1101/2021.01.11.426186

**Authors:** Mengyuan Song, Feng Song, Chenxi Zhao, Yiping Hou

## Abstract

Human Y chromosome reflects the evolutionary process of males. Male lineage tracing by Y chromosome is of great use in evolutionary, forensic, and anthropological studies when male samples exist or especially when the biological sample is a mixture of male and female individuals. Identifying the male lineage based on the specific distribution of Y haplogroups narrows down the investigation scope. Integrating previously published datasets with genotypes of Y chromosome short tandem repeats (Y-STRs) and high-resolution haplogroups (122 haplogroups in total), we developed YHP (Y Haplogroup Predictor), an open-access and userfriendly software package to predict haplogroups, compare the similarity, and conduct mismatch analysis of samples with Y-STR profiles. The software is available at Github (https://github.com/cissy123/YHP-Y-Haplogroup-Predictor-).

**Author Summary:** Familial searching has been used in forensic, anthropologic, and personalized scenarios. Software packages have been developed to assist in male familial searching, such as predicting Y-SNP haplogroups by Y-STRs. However, these software packages, in general, achieve this goal with a rough resolution. In this study, we developed a software package to conduct high-resolution haplogroup inference to help familial searching and at the same time reduce the cost, since it does not require tiresome Y-SNP sequencing.

## Introduction

Human Y chromosome has its unique evolutionary pattern and thus male phylogeny can be used to trace male lineages, which is promising in evolutionary, forensic and anthropologic studies. In forensics, identifying the possible genealogy of a DNA profile in crime scene investigations based on searching from the DNA database is of great interest (1,2). Previously findings of autosomal chromosomes indicate that some forensically useful marker sets might bear substantial ancestry information (3), indicating a significant connection between genes and geography (4). Besides, potential matches for two kinds of distinct genetic markers were reported, such as Combined DNA Index System (CODIS) profile and single nucleotide polymorphism (SNP) data, making it possible to link a CODIS profile to a whole-genome SNP profile (5–7). For Y chromosomes, the correlation of surnames and male-specific region markers in Y chromosome is vital (8,9). Since surnames are arranged by male lineage in general, we wondered if there was a correlation between two kinds of Y-chromosome markers, Y-STRs and Y-SNPs (markers defining Y haplogroups), especially in haplogroup O.

Due to the low cost-effectiveness to genotype plenty of SNPs to assign haplogroups to individuals, and the link between Y-STR variability and haplogroups (10), many software or programs appeared (**Table 1**). The software named “Yleaf’ was established for Y haplogroup inference from next-generation sequencing data (11), as well as many other packages for Y-STR data (12). Similarly, algorithms have been raised to classify mtDNA haplogroups (13). Previously, machine learning methods have been largely used in biological studies. Random forest has been previously used in reconstructing invasion routes of Drosophila suzukii using a multi-locus microsatellite dataset containing 25 loci of 23 population sites (14). Support Vector Machine (SVM) was used to inference the biogeographic ancestry based on STR profiles (15). Deep neural networks were also applied in predicting geographic location using whole-genome sequence data of the organisms, achieving median test errors of 16.9km, 5.7km and 85km for three species (Plasmodium parasites, Anopheles mosquitoes, and global human populations) (16). More specifically, artificial neural networks were also used in classifying electrophoresis profiles in forensic casework (17,18). Here in this study, we used machine learning to predict Y haplogroups to a fine resolution based on Y-STRs.

**Table 1.**
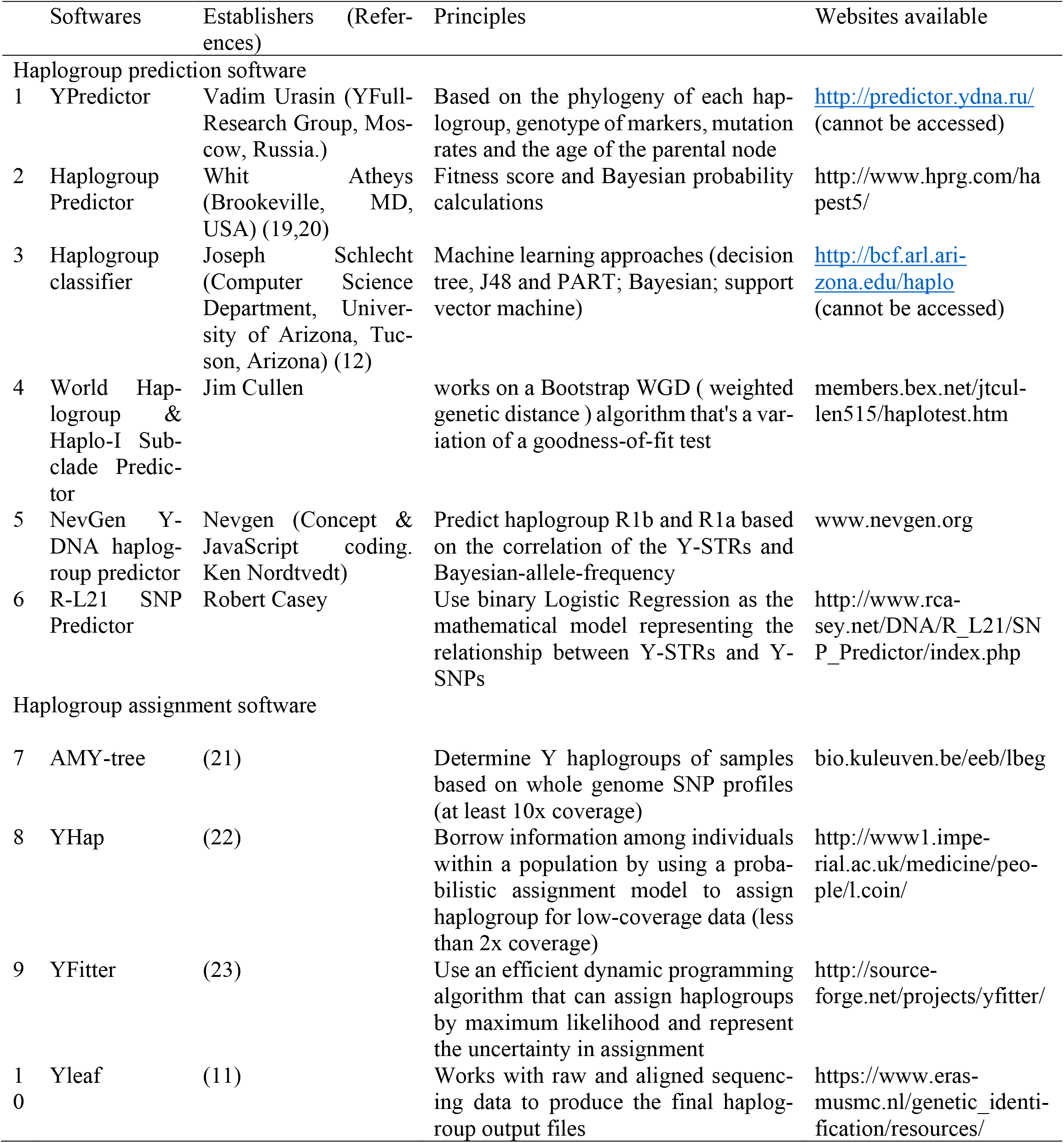
Summary of previous softwares.

## Results and Discussion

Here we present YHP (Y Haplogroup Predictor), based on machine learning algorithms, written in Java, a user-friendly public software package to predict Y haplogroups based on Y-STRs. The prediction accuracy was shown in **FIG. 1A**. Haplogroup information of database samples used to train the algorithms was illustrated in **FIG. 1B** (detailed haplogroup information is in **Supplementary table 1**). The three functions of YHP are shown in **FIG. 1C**.

**Fig 1.**
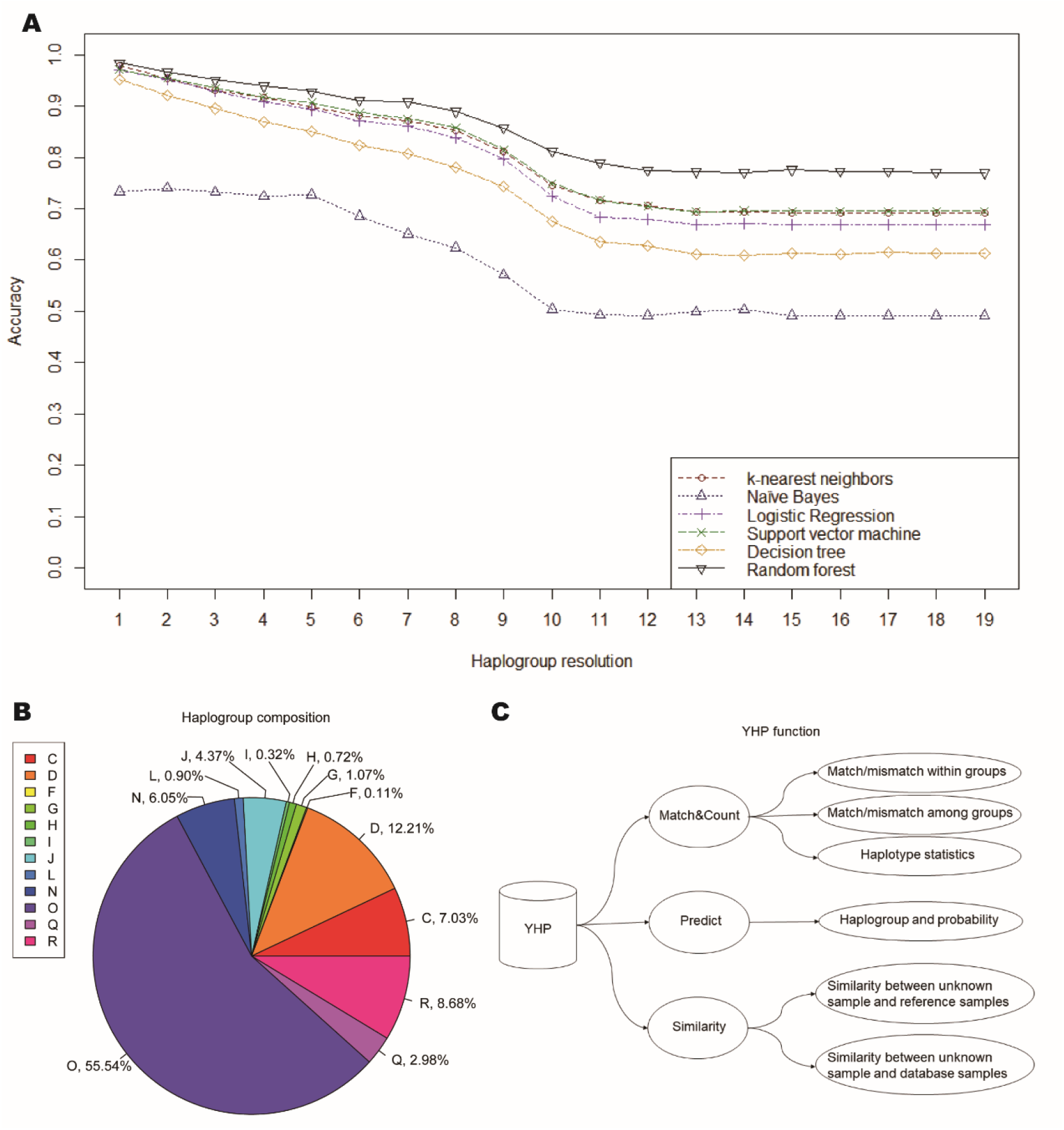
(A) Prediction accuracy of different models under different haplogroup resolution (number 1-19 means the length of the haplogroup name). (B) Haplogroup composition of the database. (C) Three main functions of YHP and the expected results.

Of the six algorithms, random forest achieved the highest accuracy (both in the terminal and basal haplogroup: 0.770 and 0.984, respectively). Prediction accuracy was defined by the number of samples correctly predicted dividing the total sample size of the training datasets and was shown in **Table 2**. Except for haplogroup prediction, we conducted population and region prediction. However, the accuracy is lower when predicting for population and region (**Table 2**). More specifically, the accuracy for each haplogroup in random forest was displayed in **FIG. 2**.

**Fig 2.**
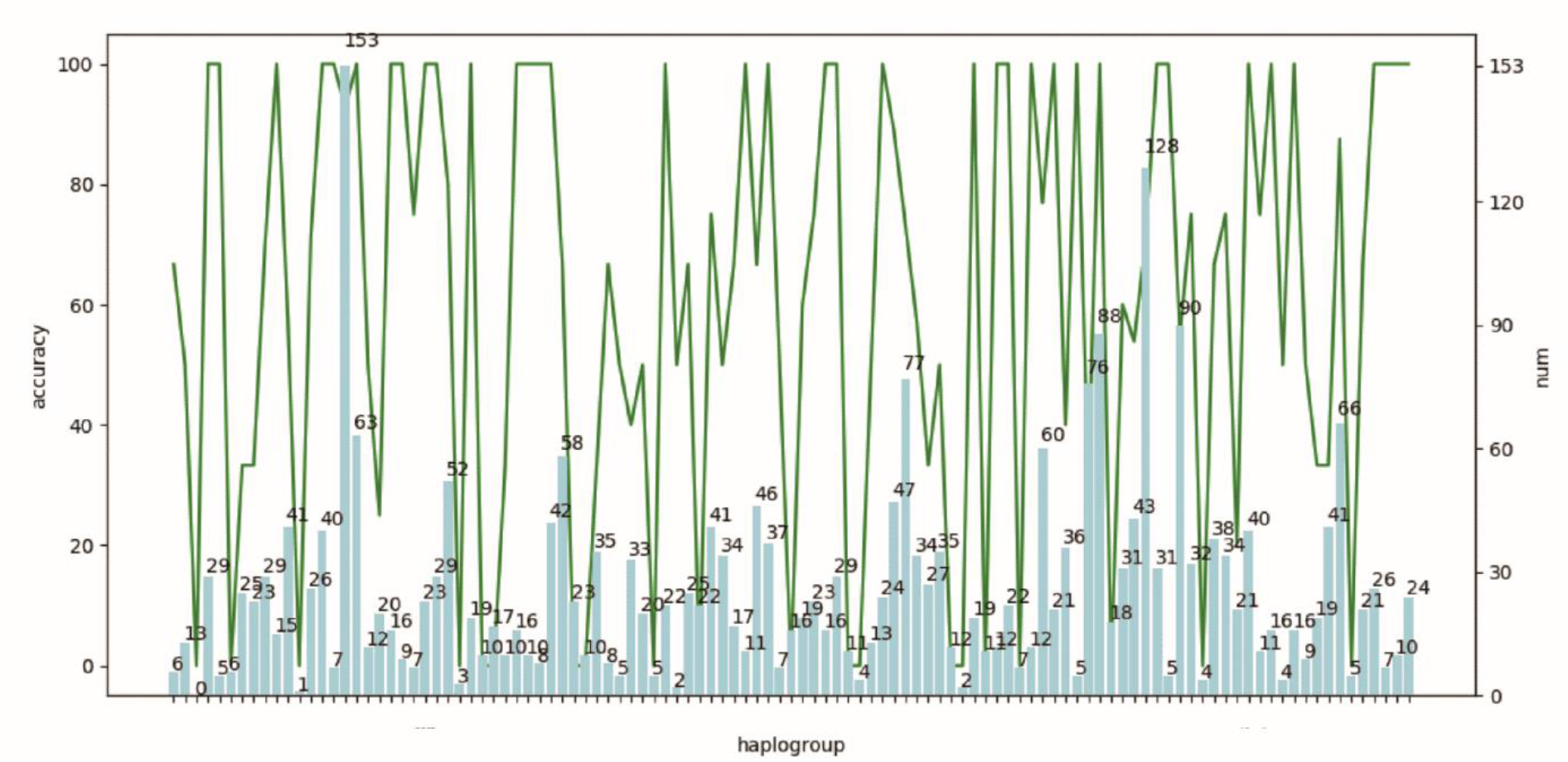
Prediction accuracy for each haplogroup in random forest. The number in each bar indicates the sample size in each haplogroup. The haplogroup information in x-axis can be obtained upon request.

**Table 2.** Prediction accuracy of the six acquired models while predicting the sample to haplogroup, population or region.

| Methods | Accuracy for haplogroup | population | region |
| --- | --- | --- | --- |
| k-nearest neighbors | 0.691 | 0.671 | 0.235 |
| Naïve Bayes* | 0.735 | 0.568 | 0.121 |
| Logistic Regression* | 0.736 | 0.627 | 0.136 |
| Support vector machine* | 0.738 | 0.721 | 0.209 |
| Decision tree* | 0.659 | 0.623 | 0.189 |
| Random forest | 0.770 | 0.752 | 0.255 |

\*The methods are optimized by linear discriminant analysis (LDA).

The use of YHP was previously validated in a real case (24), population samples (25), and another case (seen in **Supplementary figure 1–3 and Supplementary table 2–4, Supplementary Material I**), all of them with validated Y-STR and Y-SNP genotypes. This was achieved by the first and second function, “Predict” and “Similarity”.

The third function, “Match&Count” serves when there is an unknown sample (e.g., from the real crime scenes or anthropological sites) and reference samples (e.g., Y-STR profiles from the database, without Y haplogroup information), and we need to find the closest male lineage to the unknown sample to conduct familial searching (fig. 1C, YHP function; the detailed function description of YHP input files, pipeline, and output files are in **Supplementary figure 4–18, Supplementary Material II, III, and IV**). This software is also convenient for mismatch analysis within or among haplogroups and populations. The function was previously applied in a paper describing the founder effect of Li ethnic group (26), and was instructive in familiar searching. We conducted 5,966,785 times mismatch (n=3455) calculations in the software and the results were shown in Supplementary table 3 and 4. The results shows, when mismatch number is no more than two, the frequency of the sample pair belonging to the same haplogroup exceeds 97% (mismatch number=0, 100%; mismatch number=1, 99.28%; mismatch number=2, 97.16%); when mismatch step is no more than two, the frequency of the sample pair belonging to the same haplogroup exceeds 97% (mismatch number=0, 100%; mismatch number=1, 99.08%; mismatch number=2, 97.22%) (**Supplementary table 5 and 6**).

Previous relevant software or programs aim at predicting samples to haplogroup I, R, J or very basal haplogroups (seen in Figure 1 of (12)), or assign haplogroup based on high-coverage or low-coverage whole-genome sequencing or resequencing data (**Table 1**). For instance, inconsistency was reported in haplogroup prediction of a father-son pair using Whit Atheys’ haplogroup predictor (http://www.hprg.com/hapest5/hapest5b/hapest5.htm) (20,27). However, after Y-SNP testing, the father-son pair was validated in the same haplogroup O1a1a. This indicated that more accurate prediction is needed. The software YHP can effectively predict the father-son pair into haplogroup O1a1a2a1. YHP mainly focuses on haplogroup O (1919/3455, 55.54%, fig. 1B) (26,28,29) to give a high-resolution prediction result, where no previous software reached this resolution. We have extended the resolution to 122 terminal clades, and hopefully, in the future, the software can perform prediction more specifically without sacrificing too much accuracy.

Since it requires haplotypes with known haplogroups to obtain well-established models, a larger dataset needs to be generated to achieve higher accuracy. Admittedly, the prediction accuracy is not under satisfaction in the finest resolution (although in basal haplogroup prediction, the accuracy reaches 98.4%). However, the unprecedented high resolution of haplogroup makes the software valuable in differentiating close male lineages, thus narrowing down the investigative scope in forensic and anthropological events.

Although there might be a plethora of samples that only have a few Y-STR mismatches when searching the database, pinpointing samples that are probable to be the same haplogroup is largely restricted. STRs are appealing genetic materials about both population history and evolutionary process, but they are difficult to interpret due to the back mutations (30,31). Considering the low mutation rate of Y-SNPs, individuals with the same prediction results tend to be from the same male lineage. This is of tremendous use for familial searching to speed up the process of finding the perpetrator.

## Design and Implementation

### Datasets

Here we use 3455 samples with 27 Y-STRs and 137 Y-SNPs in the dataset (the haplogroup information is listed in **Supplementary table 1**), generated by capillary electrophoresis (Genetic Analyzer 3130 and 3500) and next-generation sequencing (Ion Torrent PGM) and pyrosequencing (26,28,29). The study received the approval of the Ethics Committee at the Institute of Forensic Medicine, Sichuan University (K2019018) and the data were analyzed anonymously due to privacy concerns.

### Algorithms

Supervised learning algorithms, k-nearest neighbors, Naïve Bayes, Logistic Regression, Support vector machine, Decision tree, and Random forest were used to train a model respectively. The acquired model was used to predict the test datasets. When training a model, we randomly split the data into training and test datasets to get a good representation of all data points. We split 3455 people into two disjoint subsets: a training set for learning associations between Y-STRs and Y-SNPs and a test set for assessing prediction accuracy (400 samples as test dataset and the remaining as training dataset; the training process was finished using 10 iterations). We use five-fold cross-validation with the same fraction of the full data (12%). The input and output variables are indicated as X and Y, respectively, while the value for these two variables is indicated by x and y. The input data x is indicated as:

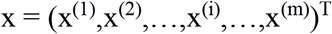

x^(i)^ is the ith locus of a single haplotype with m Y-STRs (m=27 in this study). The output data y_i_ is the haplogroup of the corresponding x_i_. The training data TR consists of pairs of input and output values, shown as:

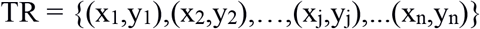

y_j_ is the haplogroup of sample j, with n being the total sample number (n=3455 in this study).

Supervised learning assumes that input and output variables X and Y are subject to the probability distribution P(X,Y), which is a probability density function. In the learning process, learning system uses the specified training dataset to learn and get a model, which is indicated as conditional probability distribution P(Y|X) or statistical decision function. In the predicting process, predicting system will give an output y_N+1_ based on the input X_N+1_ and the model:

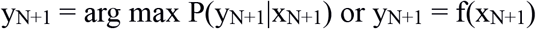

If the model has a high capability of prediction, the difference between the training data yi and the data f(x_i_) obtained from the model should be subtle enough (that means the sample is predicted to the closest haplogroup). The learning system will select the best model among all learning process to give the best prediction for the training dataset and unknown datasets.

Next, to give a rank to the reference samples evaluating the closest sample to the unknown sample, we developed similarity score using cosine distance, which is indicated as follows:

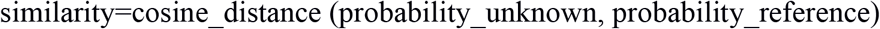

### Availability and future directions

The example data, and the code are available at Github (https://github.com/cissv123/YHP-Y-Haplogroup-Predictor-). The software YHP works under Java environment, the package of which can be downloaded from the link written in the readme file of the website.

Future directions include developing a Linux-based version and optimizing the algorithms for prediction.

## Supporting information

S1-6 Table and S1-18 Figure are compiled in the Supplementary material file (PDF).

## Author Contributions

Conceptualization: Mengyuan Song, Yiping Hou.

Data curation: Feng Song, Chenxi Zhao.

Funding acquisition: Feng Song, Yiping Hou.

Methodology: Mengyuan Song, Chenxi Zhao.

Software: Mengyuan Song, Chenxi Zhao.

Supervision: Feng Song, Yiping Hou.

Writing-original draft: Mengyuan Song.

Writing-review & editing: Mengyuan Song, Feng Song, Chenxi Zhao, Yiping Hou.

## Data Availability Statement

All relevant data are within the manuscript and its Supporting Information files.

## Supplementary material

**Supplementary table 1.** Haplogroup information of the samples and their corresponding size in each haplogroup.

| haplogroup | number of<br>haplotypes | haplogroup | number of<br>haplotypes | haplogroup | number of<br>haplotypes |
| --- | --- | --- | --- | --- | --- |
| C | 9 | N1a1a | 17 | O2a1c1a1a1b | 20 |
| C2 | 15 | N1a1a1a1a3 | 10 | O2a1c1a1b | 1 |
| C2b | 7 | N1a1a1a1a4 | 9 | O2a1c1a1c | 12 |
| C2b1a1b1 | 3 | N1a2 | 39 | O2a1c1a1d | 13 |
| C2b1a2 | 6 | N1b | 67 | O2a1c1a1e | 27 |
| C2b1a3 | 29 | N1~ | 24 | O2a2 | 8 |
| C2b1b | 6 | O1a | 10 | O2a2a | 13 |
| C2c1 | 8 | O1a1a | 15 | O2a2a1 | 73 |
| C2c1a1 | 23 | O1a1a1a | 5 | O2a2a1a1a | 23 |
| C2c1a1a1 | 26 | O1a1a1a1 | 36 | O2a2b | 42 |
| C2c1a2 | 38 | O1a1a1a1a | 14 | O2a2b1 | 1 |
| C2c1a2b | 16 | O1a1a1a1a1a | 7 | O2a2b1a1 | 6 |
| C2c1b | 41 | O1a1a1a1a1a1 | 38 | O2a2b1a1a | 84 |
| D1a1 | 6 | O1a1a1a1a1a1a | 24 | O2a2b1a1a1 | 93 |
| D1a1a1a | 3 | O1a1a1a1a1a1b | 2 | O2a2b1a1a3 | 23 |
| D1a1a1a1 | 2 | O1a1a1a1a1a1b1 | 8 | O2a2b1a1a4 | 35 |
| D1a1a1a1a | 31 | O1a1a1b | 5 | O2a2b1a1a5 | 56 |
| D1a1a1a1a~ | 9 | O1a1a1b1 | 4 | O2a2b1a1a6 | 148 |
| D1a1a1a2 | 46 | O1a1a1b2 | 27 | O2a2b1a2 | 28 |
| D1a1a1a2b | 8 | O1a1a2 | 22 | O2a2b1a2a | 6 |
| D1a2a1 | 3 | O1a1a2a1 | 35 | O2a2b1a2a1 | 99 |
| D1a2a1a~ | 2 | O1b | 5 | O2a2b1a2a1a3 | 34 |
| D1a2a1b | 171 | O1b1a1 | 35 | O2a2b1a2a1a3b1 | 5 |
| D1a2a1b1 | 6 | O1b1a1a | 6 | O2a2b1a2a1a3b2 | 39 |
| D1a2a1b1a | 66 | O1b1a1a1a | 19 | O2a2b1a2a1a3b2b2 | 38 |
| D1a2a1b2 | 15 | O1b1a1a1a1a | 6 | Q | 25 |
| D1a2a1b3 | 3 | O1b1a1a1a1a1 | 43 | Q* | 43 |
| D1a2a1b~ | 7 | O1b1a1a1a1a1b | 1 | Q1a2 | 15 |
| DE | 24 | O1b1a1a1a1a1b1 | 6 | Q1b | 18 |
| F2 | 4 | O1b1a1a1a1a2 | 19 | R | 1 |
| G | 7 | O1b1a1a1a1a1b | 25 | R1a1a | 6 |
| G2a | 8 | O1b1a1a1a1a1b1 | 22 | R1a1a1b1a1 | 6 |
| G2a2b | 1 | O1b1a1a1b | 16 | R1a1a1b1a2 | 4 |
| G2a2b2a | 3 | O1b1a2a | 29 | R1a1a1b2 | 15 |
| G2a2b2a1 | 15 | O1b1a2b | 12 | R1a1a1b2a | 11 |
| H1a | 11 | O1b1a2c | 5 | R1a1a1b2a1a | 1 |
| H1a1a | 8 | O1b2 | 19 | R1a1a1b2a1a1a | 22 |
| H1a2a | 1 | O2 | 24 | R1a1a1b2a2 | 48 |
| I | 11 | O2a1 | 47 | R1a1a1b2a2a | 70 |
| J1 | 29 | O2a1c | 90 | R1a1a1b2a2b | 4 |
| J2 | 18 | O2a1c1a | 48 | R1a1a1b2a2b1 | 6 |
| J2a | 35 | O2a1c1a1 | 31 | R1b | 22 |
| J2a1 | 54 | O2a1c1a1a1 | 14 | R1b1a1 | 26 |
| J2a1a | 7 | O2a1c1a1a1a | 32 | R1b1a1a2 | 8 |
| J2a1b | 3 | O2a1c1a1a1a1 | 14 | R2 | 13 |
| L | 19 | O2a1c1a1a1a1a | 10 | R2a | 23 |
| LT | 10 | O2a1c1a1a1a1a1a1 | 1 |  |  |
| N | 2 | O2a1c1a1a1a1a1a1a1a | 2 |  |  |
| N1 | 17 | O2a1c1a1a1a1a1a1a1b | 4 |  |  |
| N1a | 13 | O2a1c1a1a1a2 | 3 |  |  |

### Supplementary Material I: Application in another real case

There was a target sample with Y-STR profile (unknown sample) and 25 reference samples that have the least mismatch with the unknown sample, retrieved from local Y-STR database (**Supplementary figure 1**).

**Supplementary figure 1.**
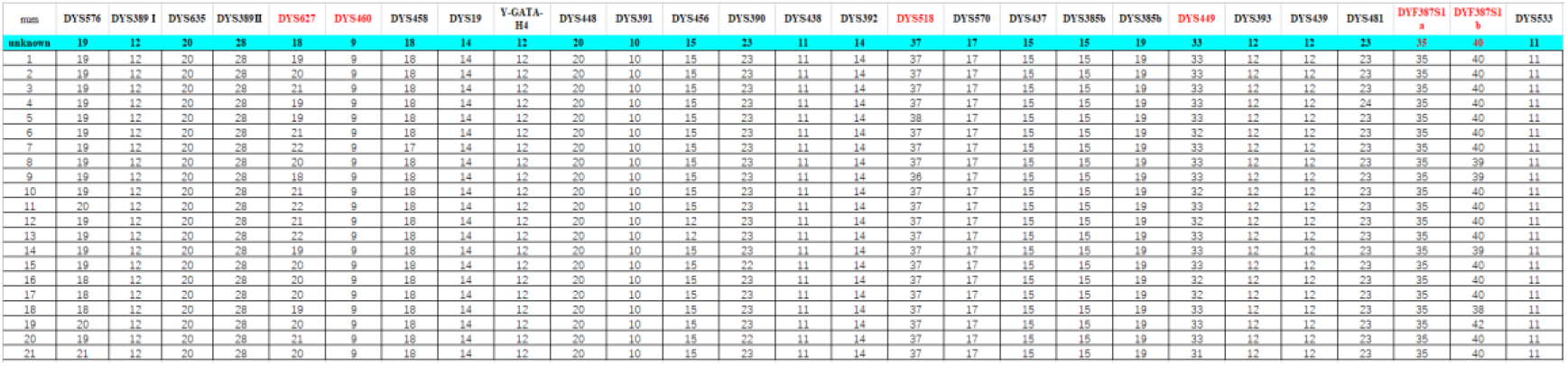
Haplotypes of a target sample and reference samples

Here questions came. Which samples are from the same male lineage as the unknown sample? What is the ranking of the reference samples according to the closeness to the unknown sample?

We used the software to compare the similarity of the unknown sample and the reference samples. Because of the different principles behind the algorithms, we calculated similarity score between the unknown sample and 25 reference samples and concluded that reference sample 23 is the closest to the unknown sample. The steps are as follows.

First, we calculated the similarity score of these reference samples to the unknown samples in three models (**Supplementary table 2, Supplementary figure 2**):

**Supplementary table 2.** Similarity score of these reference samples to the unknown samples in three models.

| Reference<br>sample | Logistic<br>Regression | SVM | Random<br>Forest |
| --- | --- | --- | --- |
| 1 | 0.992562 | 0.996092 | 0.999531 |
| 2 | 0.974887 | 0.9937 | 0.996514 |
| 3 | 0.954657 | 0.995065 | 0.996975 |
| 4 | 0.993527 | 0.998197 | 0.999751 |
| 5 | 0.983984 | 0.996739 | 0.999284 |
| 6 | 0.953399 | 0.994144 | 0.996468 |
| 7 | 0.927685 | 0.98888 | 0.998367 |
| 8 | 0.976826 | 0.989644 | 0.978107 |
| 9 | 0.98839 | 0.998236 | 0.978932 |
| 10 | 0.953399 | 0.994144 | 0.996468 |
| 11 | 0.946554 | 0.970426 | 0.993781 |
| 12 | 0.955503 | 0.937706 | 0.995788 |
| 13 | 0.938212 | 0.859445 | 0.996212 |
| 14 | 0.991501 | 0.991767 | 0.977985 |
| 15 | 0.977748 | 0.992833 | 0.996146 |
| 16 | 0.958325 | 0.988286 | 0.997434 |
| 17 | 0.958325 | 0.988286 | 0.997434 |
| 18 | 0.978671 | 0.98716 | 0.976332 |
| 19 | 0.979954 | 0.82383 | 0.994687 |
| 20 | 0.96622 | 0.995185 | 0.9964 |
| 21 | 0.972685 | 0.915663 | 0.977023 |
| 22 | 0.995496 | 0.999301 | 0.999899 |
| 23 | 0.999982 | 1 | 1 |
| 24 | 0.991264 | 0.997862 | 0.999859 |
| 25 | 0.988588 | 0.999885 | 0.999936 |

**Supplementary figure 2.**
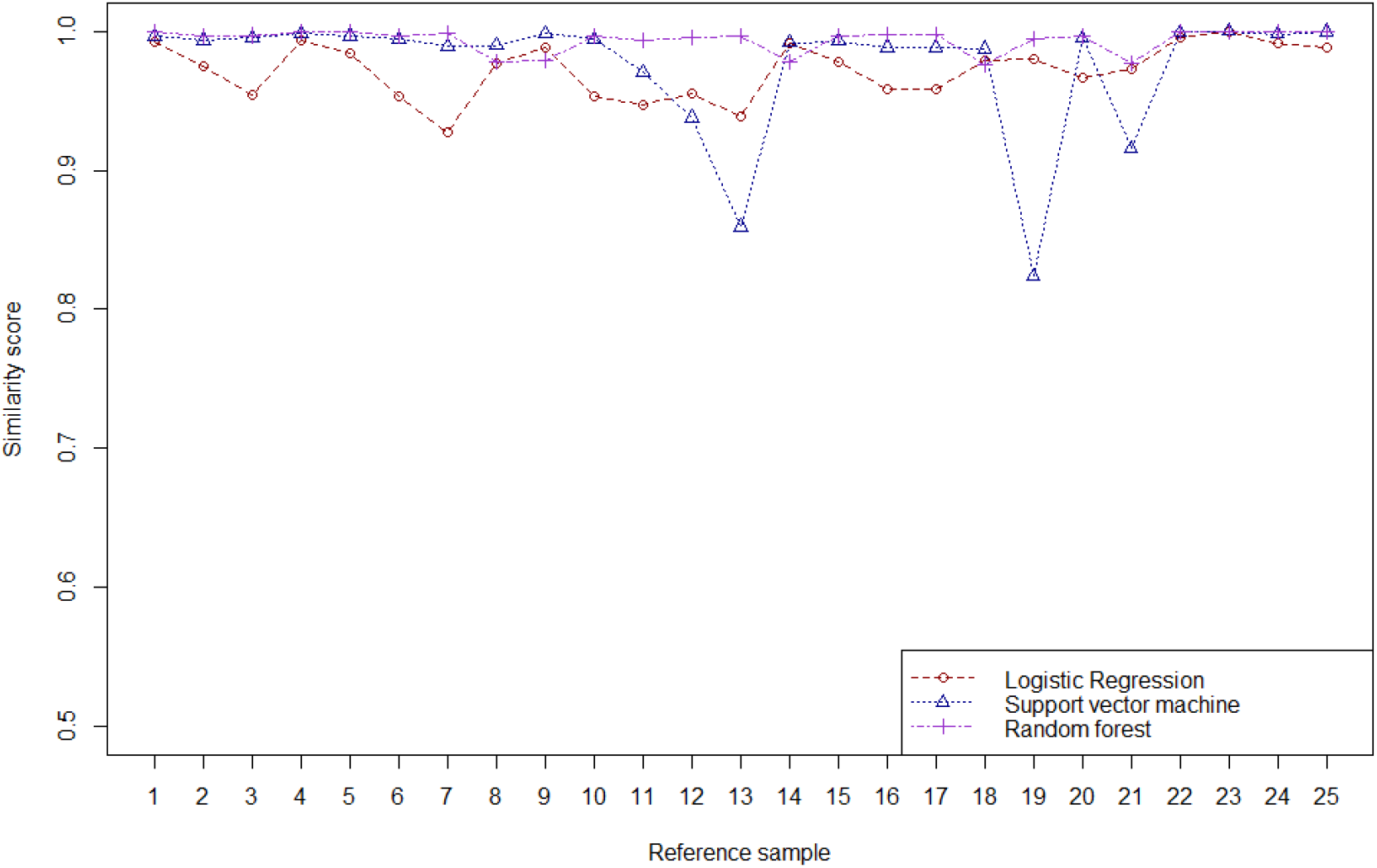
The line plot of the similarity of the unknown sample and the reference samples

Then, based on the accuracy in **Table 1**, we calculated the weighted similarity score (**Supplementary table 3, Supplementary figure 3**):

**Supplementary table 3.** Weighted similarity score by the accuracy of different models.

| Reference sample | Logistic Regression (weighted) | SVM (weighted) | Random Forest (weighted) |
| --- | --- | --- | --- |
| 1 | 0.664024 | 0.692284 | 0.769639 |
| 2 | 0.652199 | 0.690621 | 0.767316 |
| 3 | 0.638666 | 0.69157 | 0.767671 |
| 4 | 0.664669 | 0.693747 | 0.769808 |
| 5 | 0.658285 | 0.692733 | 0.769448 |
| 6 | 0.637824 | 0.69093 | 0.767281 |
| 7 | 0.620621 | 0.687271 | 0.768742 |
| 8 | 0.653497 | 0.687803 | 0.753142 |
| 9 | 0.661233 | 0.693774 | 0.753777 |
| 10 | 0.637824 | 0.69093 | 0.767281 |
| 11 | 0.633245 | 0.674446 | 0.765211 |
| 12 | 0.639231 | 0.651705 | 0.766756 |
| 13 | 0.627664 | 0.597314 | 0.767083 |
| 14 | 0.663314 | 0.689278 | 0.753049 |
| 15 | 0.654114 | 0.690019 | 0.767032 |
| 16 | 0.641119 | 0.686859 | 0.768024 |
| 17 | 0.641119 | 0.686859 | 0.768024 |
| 18 | 0.654731 | 0.686076 | 0.751776 |
| 19 | 0.655589 | 0.572562 | 0.765909 |
| 20 | 0.646401 | 0.691653 | 0.767228 |
| 21 | 0.650726 | 0.636386 | 0.752308 |
| 22 | 0.665986 | 0.694515 | 0.769922 |
| 23 | 0.668988 | 0.695 | 0.77 |
| 24 | 0.663156 | 0.693514 | 0.769891 |
| 25 | 0.661366 | 0.69492 | 0.769951 |

**Supplementary figure 3.**
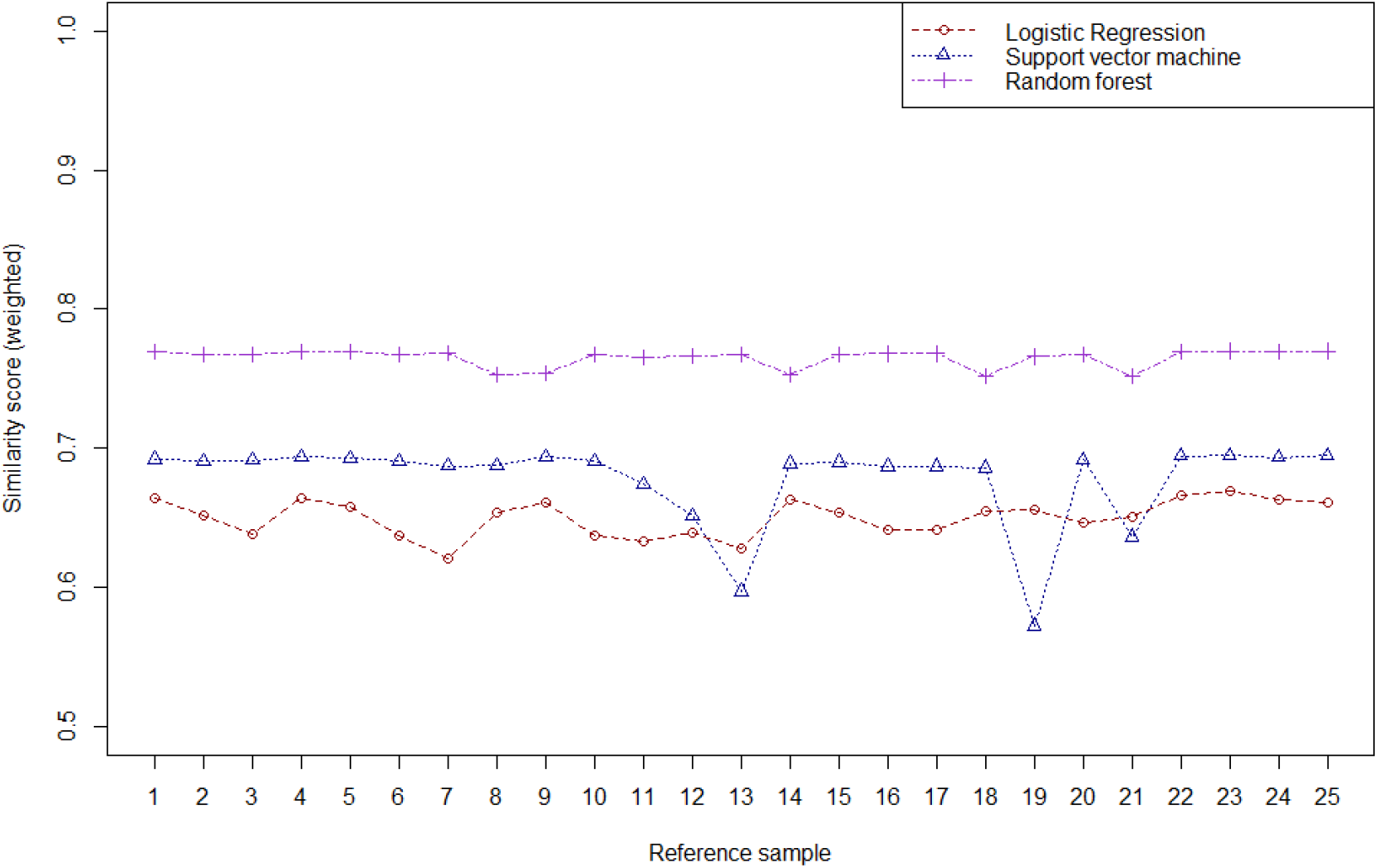
The line plot of the weighted similarity score of the unknown sample and the reference samples:

Finally, the ranking of the reference samples is based on the mean value of three weighted scores (**Supplementary table 4**):

**Supplementary table 4.** The ranking of the closeness to the unknown sample.

| Reference sample | weighted score |
| --- | --- |
| 23 | 0.711329168 |
| 22 | 0.710141087 |
| 4 | 0.709408108 |
| 24 | 0.708853689 |
| 25 | 0.708745317 |
| 1 | 0.708648946 |
| 5 | 0.706822429 |
| 15 | 0.703721642 |
| 2 | 0.703378919 |
| 9 | 0.702928022 |
| 14 | 0.701880327 |
| 20 | 0.701760818 |
| 3 | 0.699302072 |
| 6 | 0.698678134 |
| 10 | 0.698678134 |
| 16 | 0.698667317 |
| 17 | 0.698667317 |
| 8 | 0.698147231 |
| 18 | 0.697527623 |
| 7 | 0.692211636 |
| 11 | 0.690967345 |
| 12 | 0.685897656 |
| 21 | 0.679806661 |
| 19 | 0.66468681 |
| 13 | 0.66402034 |

### Supplementary Material II: YHP input files

#### II-1 “Match&Count” for mismatch analysis

Input file includes sample ID (necessary), population, haplogroup and Y-STR (necessary) genotypes (**Supplementary figure 4**).

**Supplementary figure 4.**
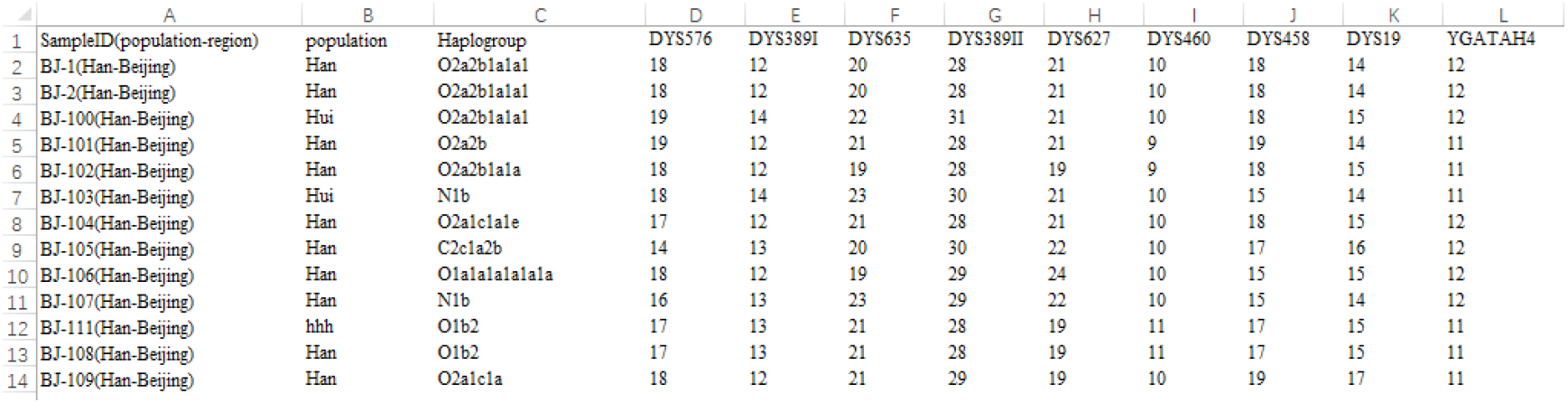
Example file: input1

#### II-2 “Predict” for haplogroup prediction

Input file includes sample ID and Y-STR genotypes (single sample: **Supplementary figure 5**; multiple sample: **Supplementary figure 6**).

#### II-2-1 Single sample mode

**Supplementary figure 5.**
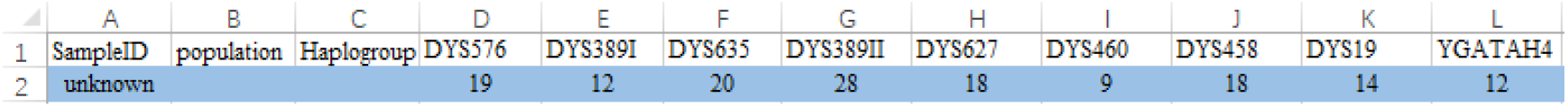
Example file: input2

#### II-2-2 Multiple sample mode

**Supplementary figure 6.**
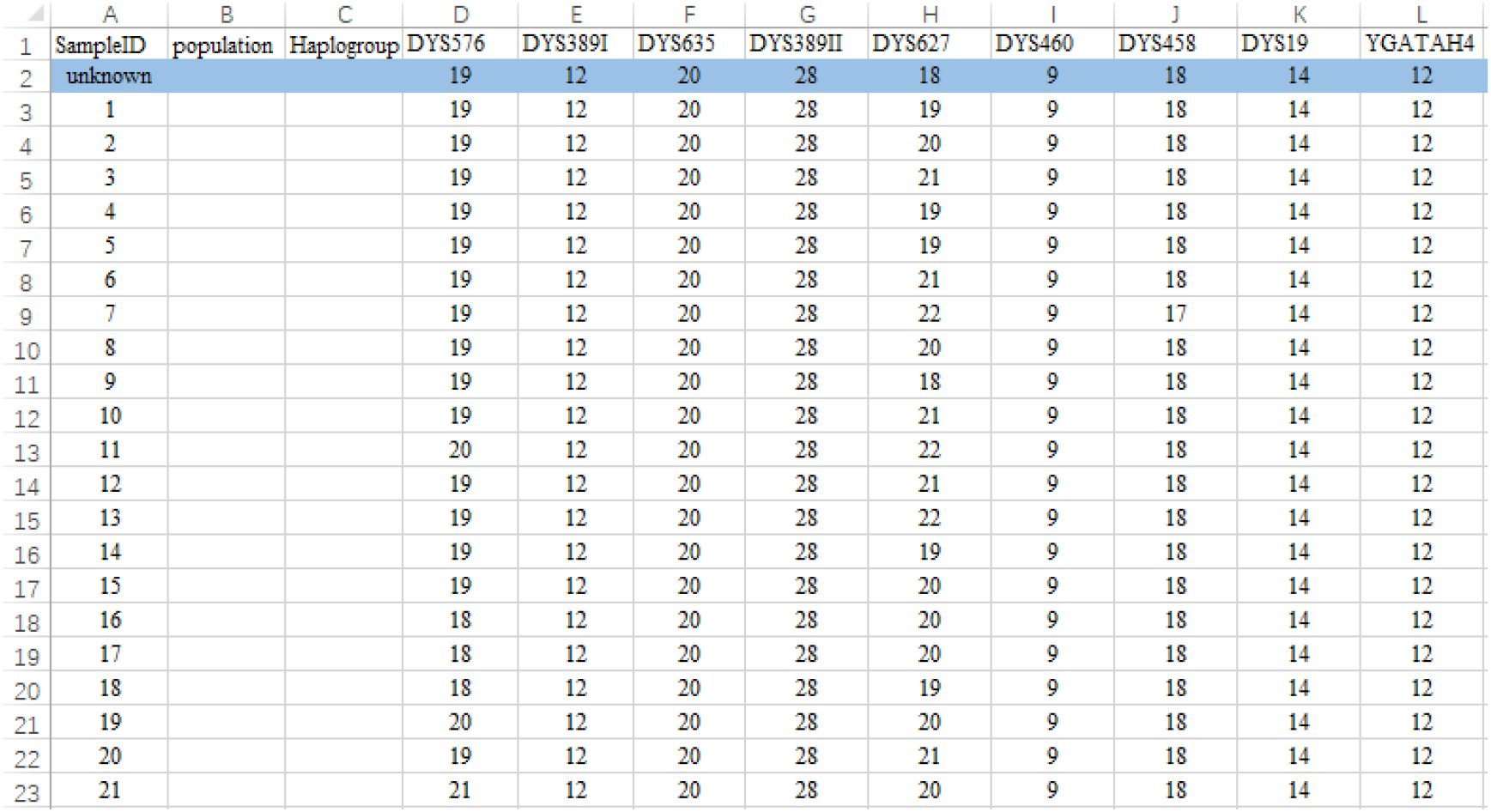
Example file: input3. The line in blue background is the unknown sample and the lines below that are reference samples.

#### II-3 “Similarity” for similarity scoring

Input file includes sample ID and Y-STR genotypes of the unknown sample and the reference samples (same as II-2-2). When there is no reference sample, the output file is mismatch result of the unknown sample and all samples in the database.

#### Supplementary Material III: YHP pipeline

For the first function Match&Count, the interface is as follow (**Supplementary figure 7 and 8**). Click the buttons to choose the input file and comparison mode.

**Supplementary figure 7.**
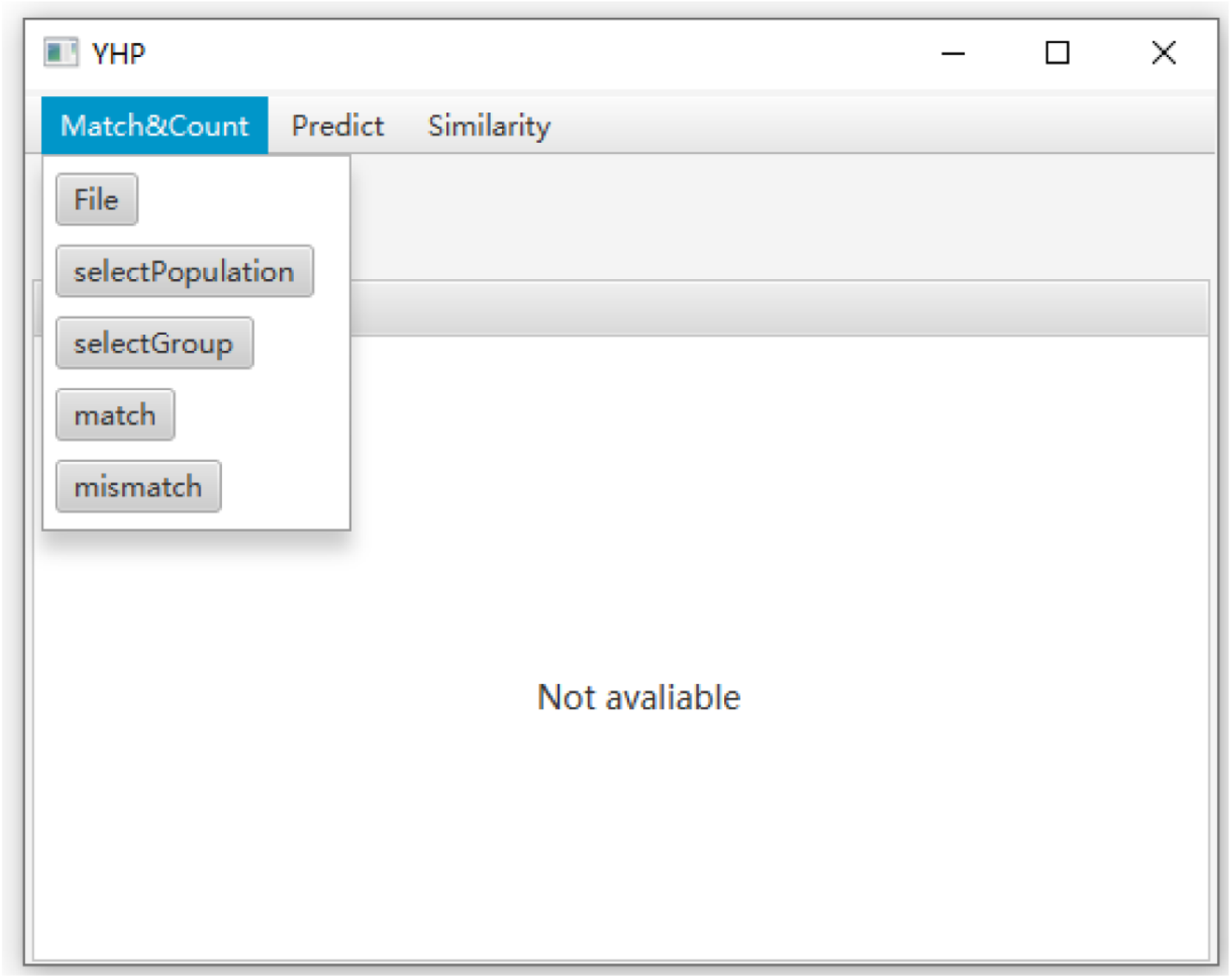
Software interface for Match&Count.

**Supplementary figure 8.**
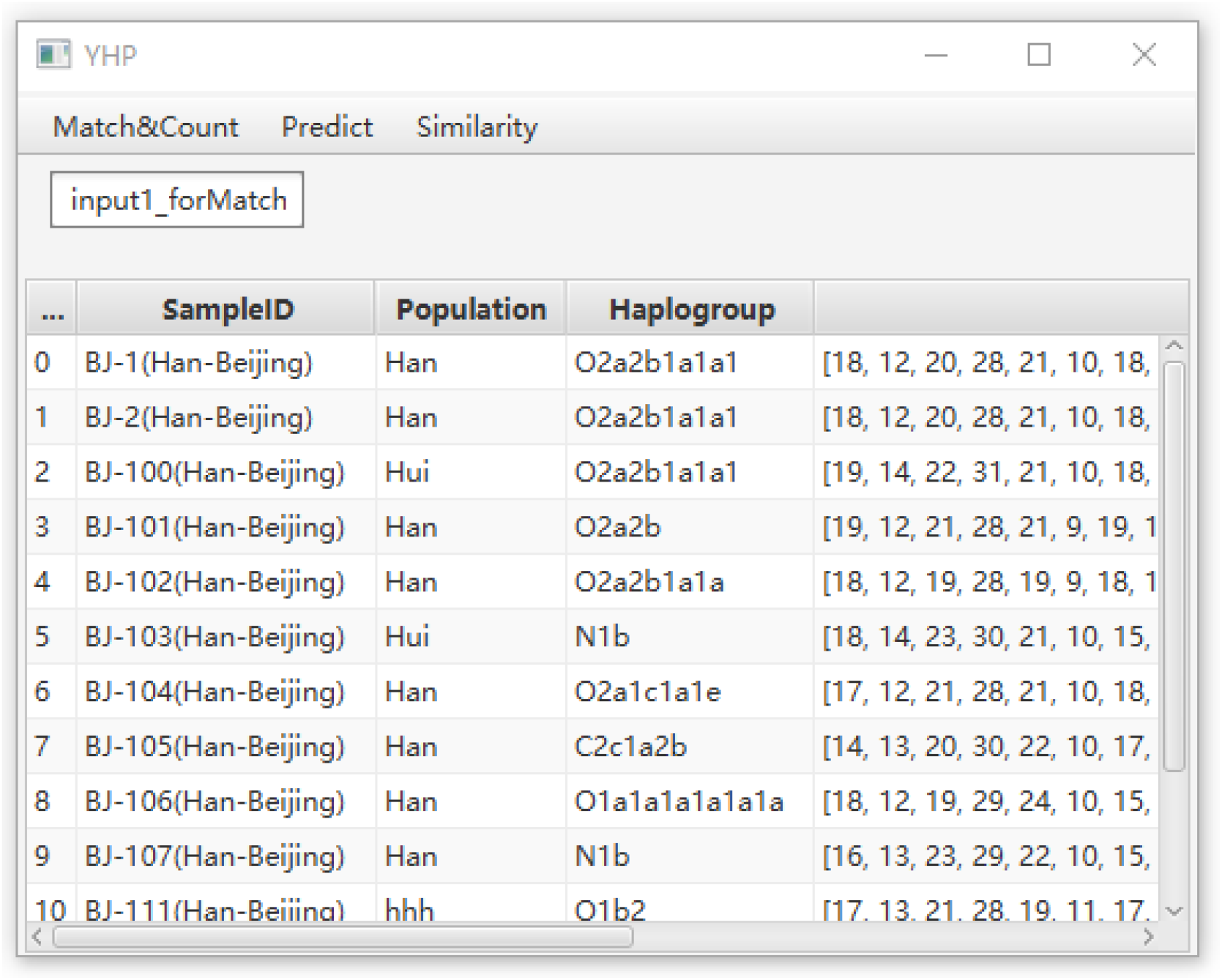
Software interface after the input file was chosen.

For the second function Predict, the interface is as follow (**Supplementary figure 9 and 10**). In this step, training data is changeable, whether using default dataset (generated in our lab as described above) or customized data (haplotypes with haplogroup information). If one uses customized data, the number of Y-STR loci is flexible, not having to be 27, but the test data should be consistent with the training data. After selection test data, click “Train” first, and then “Test”.

**Supplementary figure 9.**
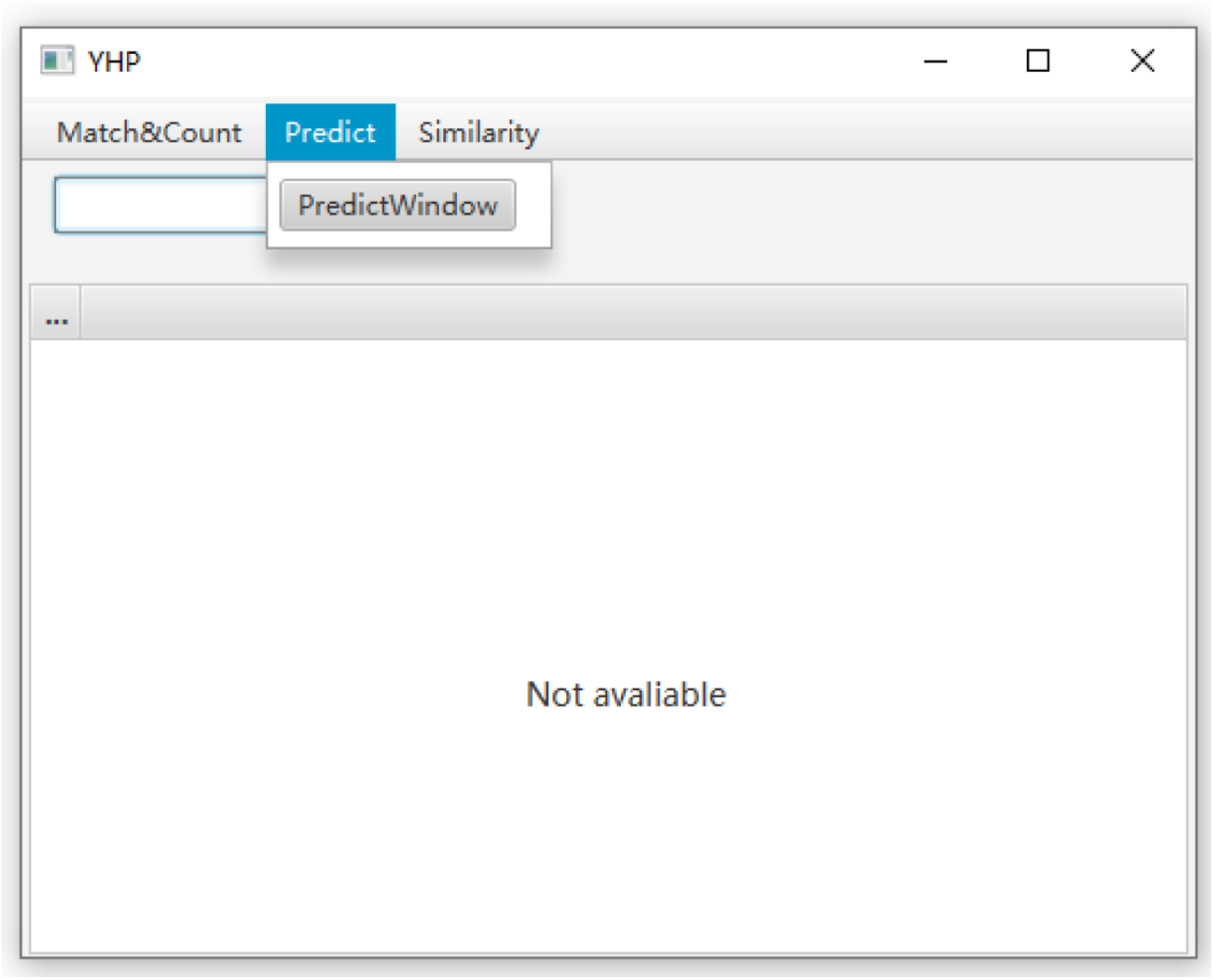
Software interface for Predict.

**Supplementary figure 10.**
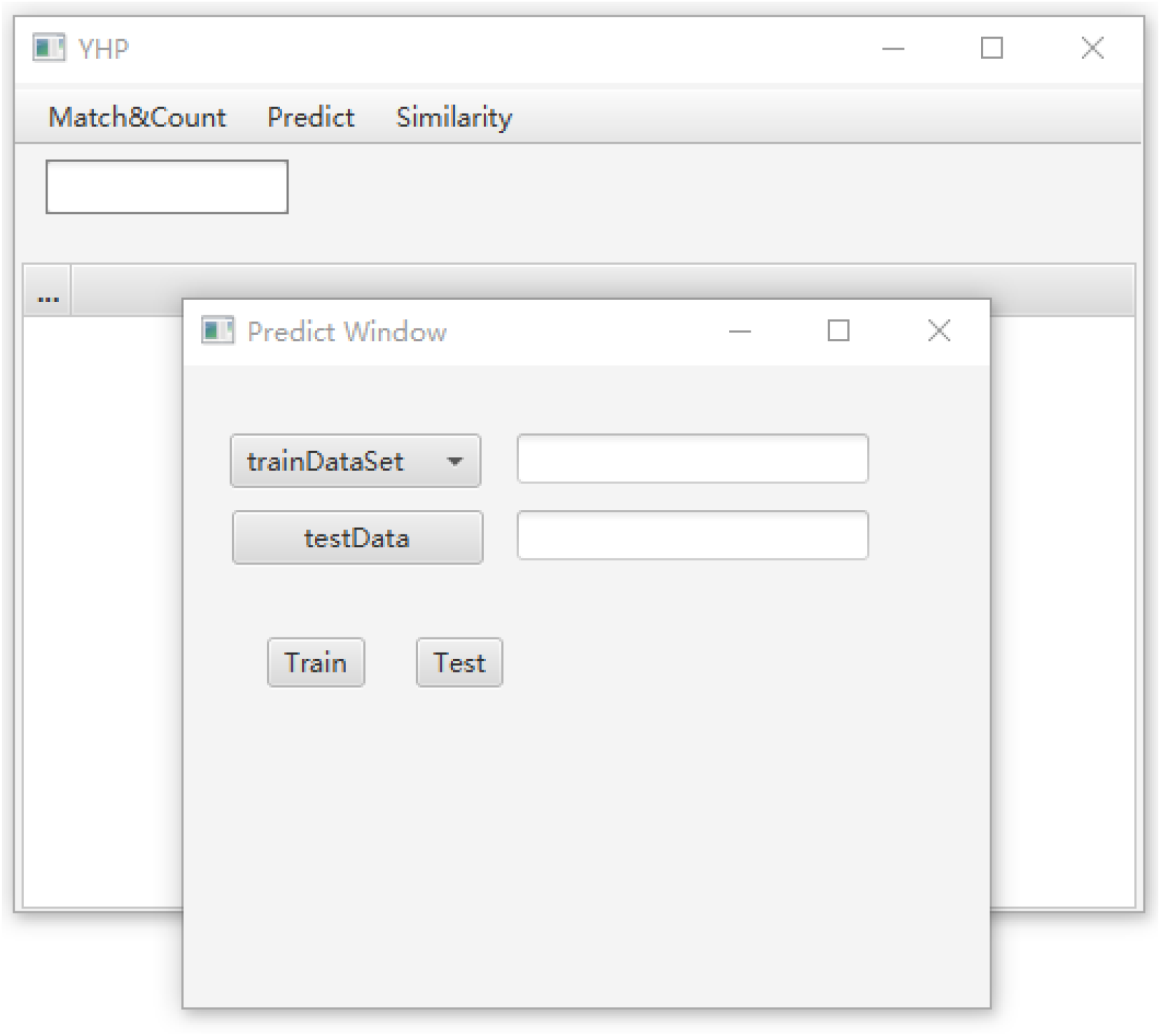
Software interface when choosing training data and test data.

For the third function Similarity, the interface is as follow (**Supplementary figure 11, 12 and 13**). There are two comparison mode, “withDatabase” and “withinSamples”, which require different input files as illustrated previously in the input section.

**Supplementary figure 11.**
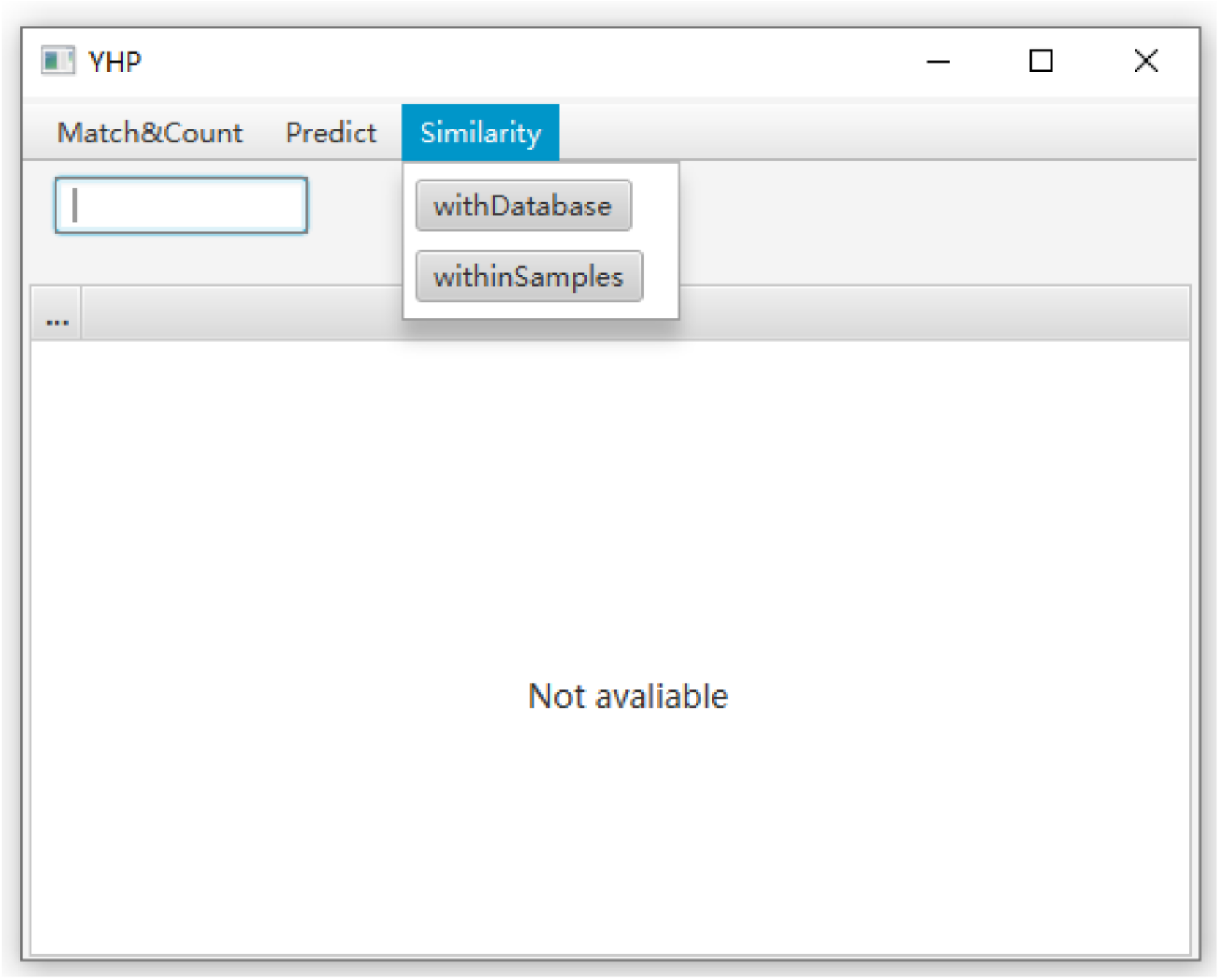
Software interface for Similarity.

**Supplementary figure 12.**
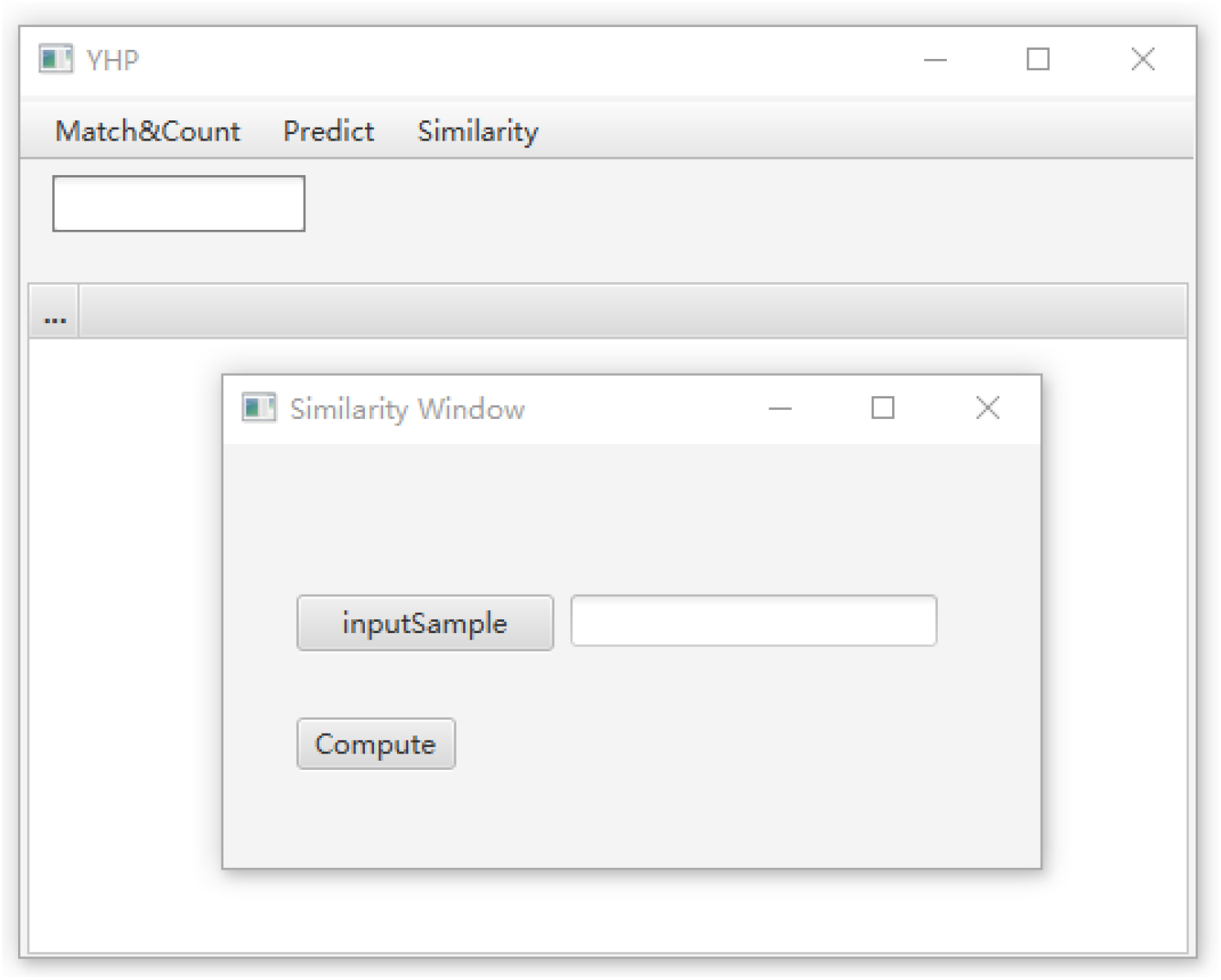
Software interface for Similarity in “withDatabase” mode.

**Supplementary figure 13.**
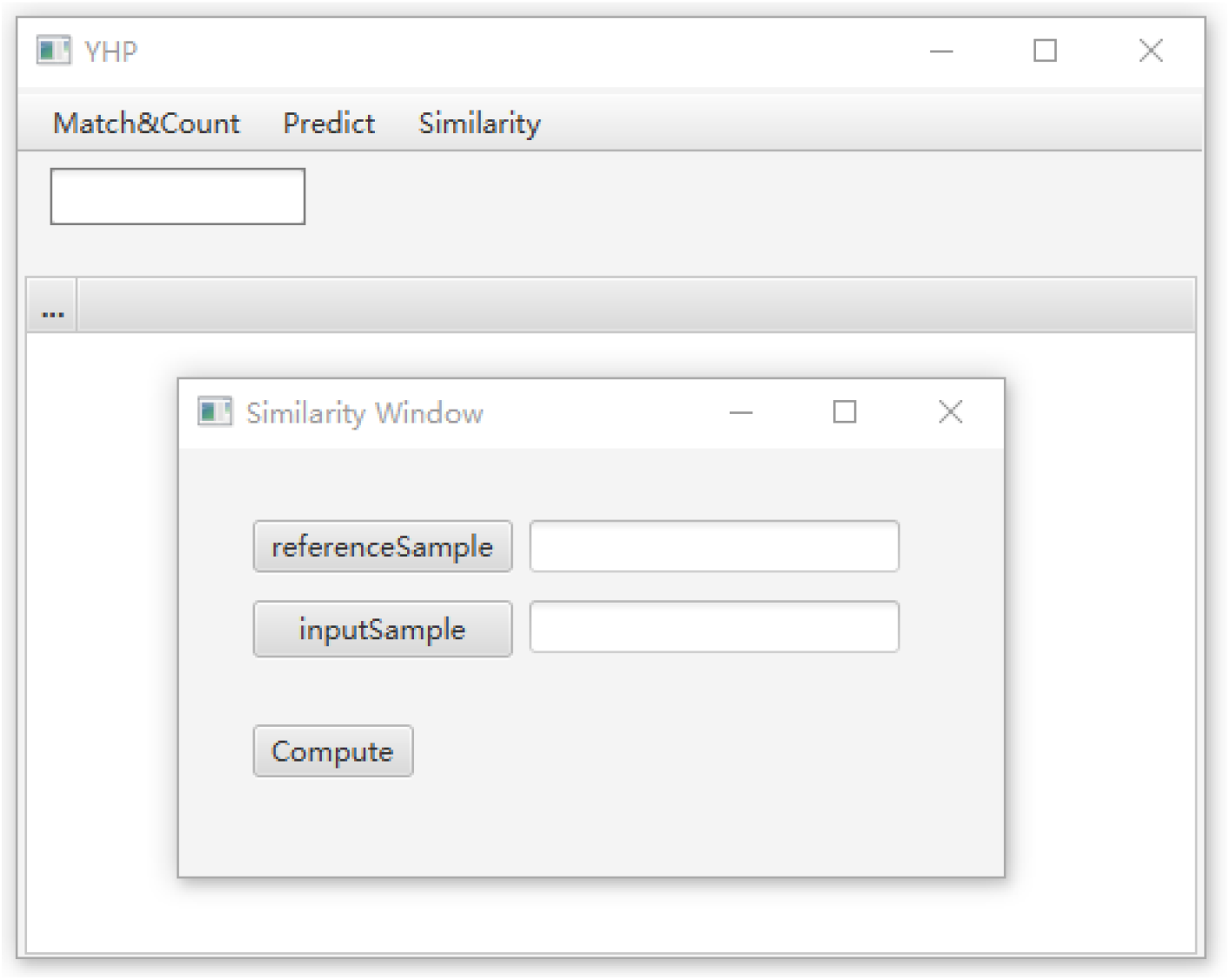
Software interface for Similarity in “withinSamples” mode

### Supplementary Material IV: Output results

The output files can be saved manually or automatically in file container “output”.

#### IV-1 Match&Count

The file can be saved in the main window after mismatch analysis. The output result includes match/mismatch number, step, ratio, and mismatch detail (**Supplementary figure 14** is one of the output results in this function).

**Supplementary figure 14.**
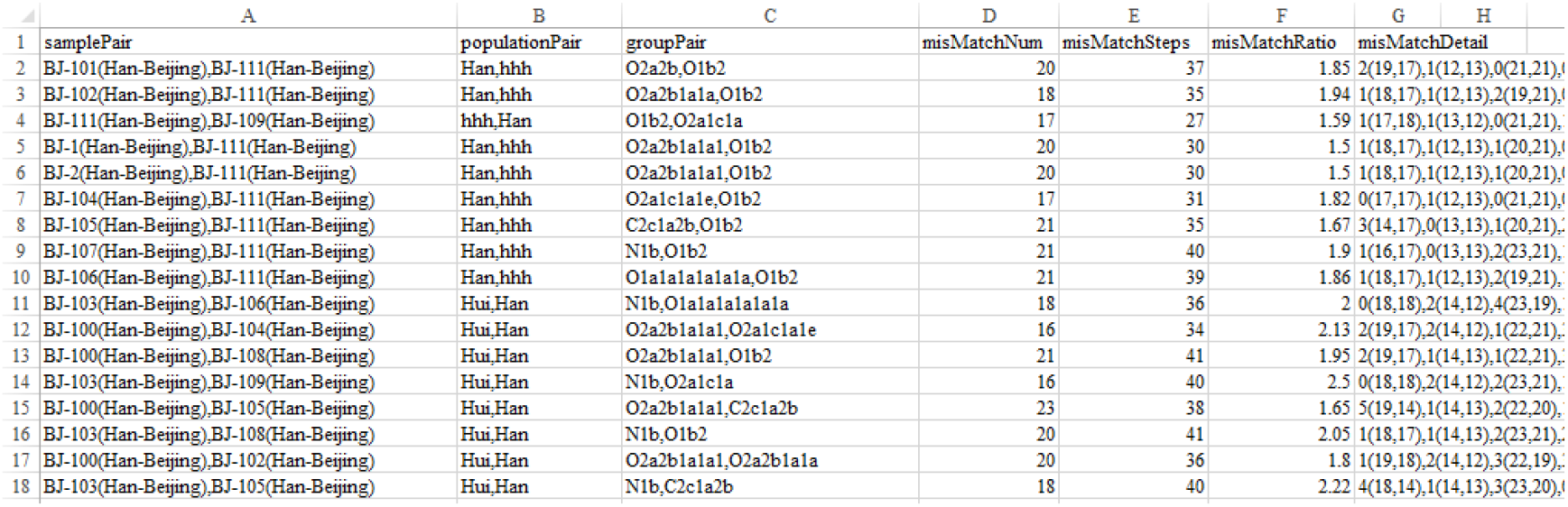
Output file for Match&Count.

#### IV-2 Predict

The predicting result (single sample: **Supplementary figure 15**; multiple sample: **Supplementary figure 16**) is saved automatically in file container “output”.

**Supplementary figure 15.**
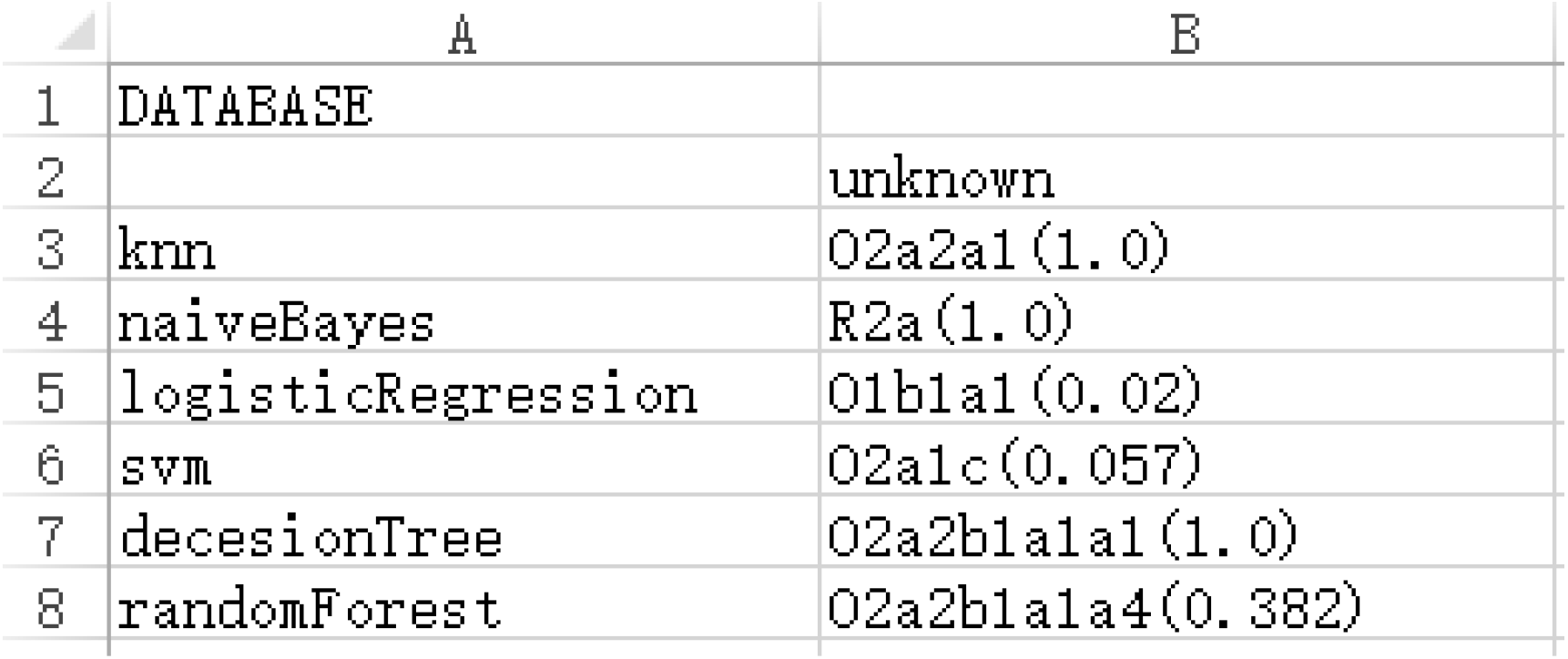
Single sample prediction result.

**Supplementary figure 16.**
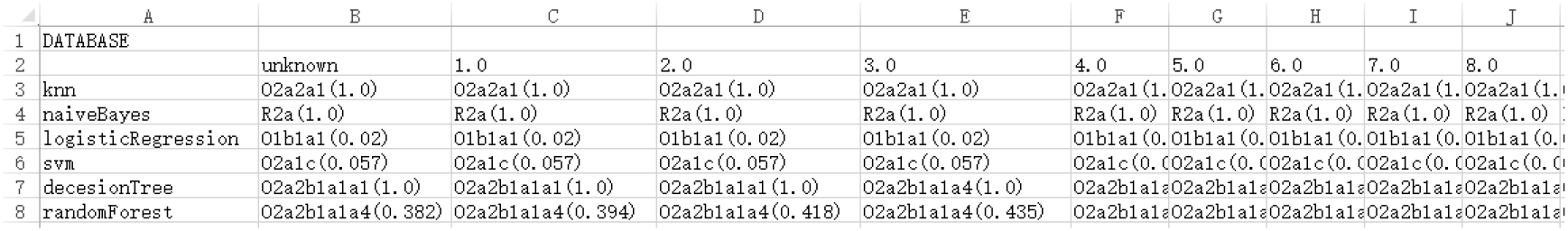
Multiple sample prediction result.

#### IV-3. Similarity

The similarity result (withDatabase: **Supplementary figure 17**; withinsamples: **Supplementary figure 18**) is saved automatically in file container “output”.

**Supplementary figure 17.**
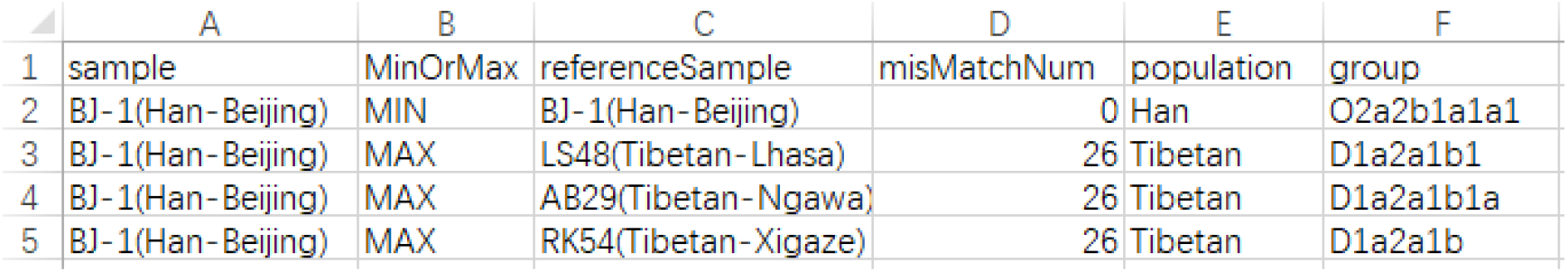
Similarity result in “withDatabase”. MIN indicates the closest sample between the target sample and samples in the database; MAX indicates the least closest sample.

**Supplementary figure 18.**
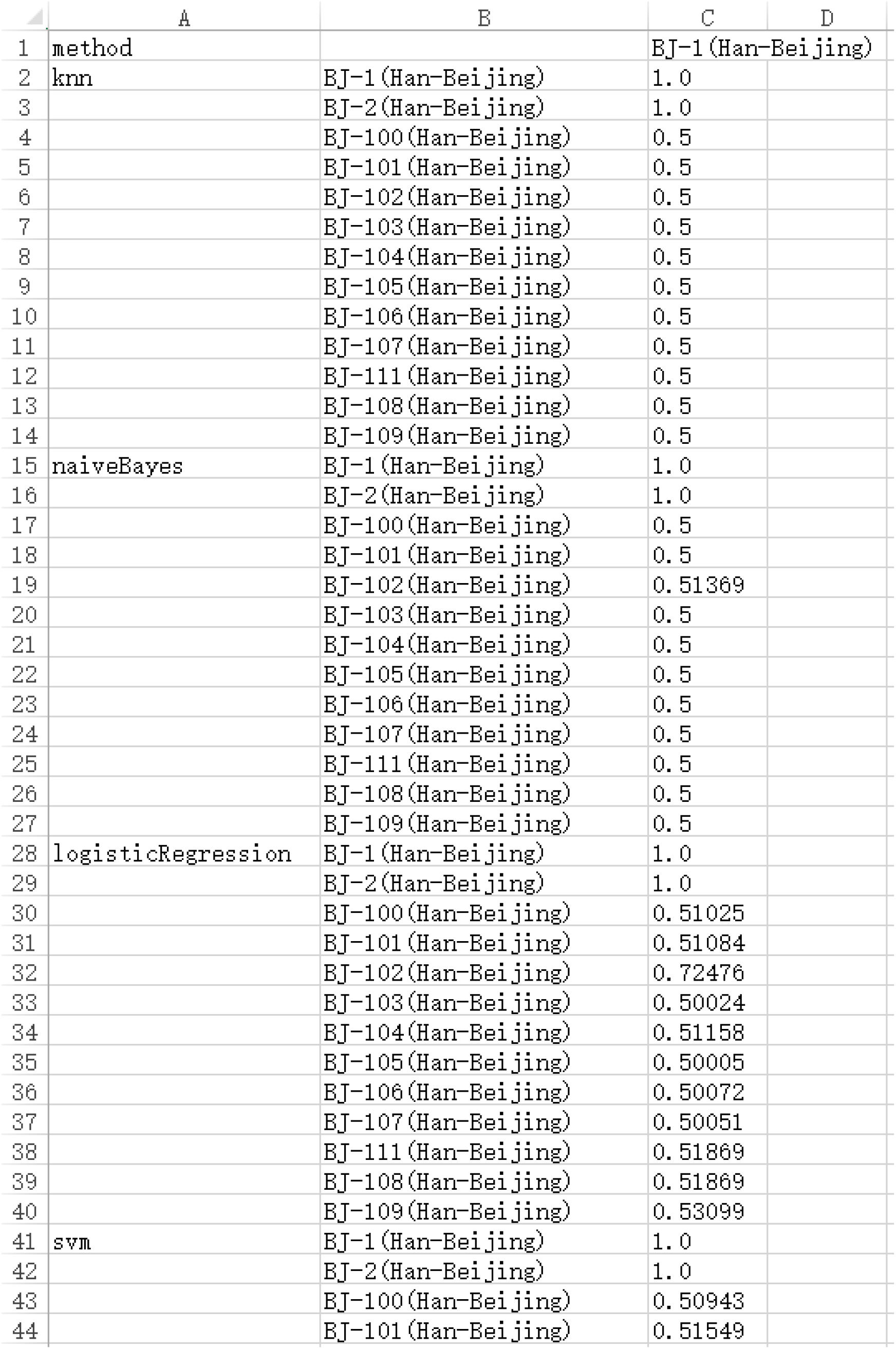
Similarity result in “withinSamples”.

The newest accessed version is up to December 7, 2020. The software will be regularly updated and Linux-based version will be released soon.

**Supplementary table 5.** Indications from mismatch number results (mismatch number is the total number of different alleles). Sample pairs are the number of pairs in the corresponding mismatch number.

| Mismatch number | Sample pairs | Pairs belonging to the same haplogroup | Pairs belonging to different haplogroups | Percentage of pairs belonging to different haplogroups |
| --- | --- | --- | --- | --- |
| 0 | 89 | 89 | 0 | 0 |
| 1 | 116 | 114 | 2 | 1.724% |
| 2 | 211 | 205 | 6 | 2.844% |
| 3 | 449 | 428 | 21 | 4.677% |
| 4 | 820 | 751 | 69 | 8.415% |
| 5 | 1565 | 1300 | 265 | 16.932% |
| 6 | 2462 | 1869 | 593 | 24.086% |
| 7 | 4221 | 2721 | 1509 | 35.750% |

**Supplementary table 6.**
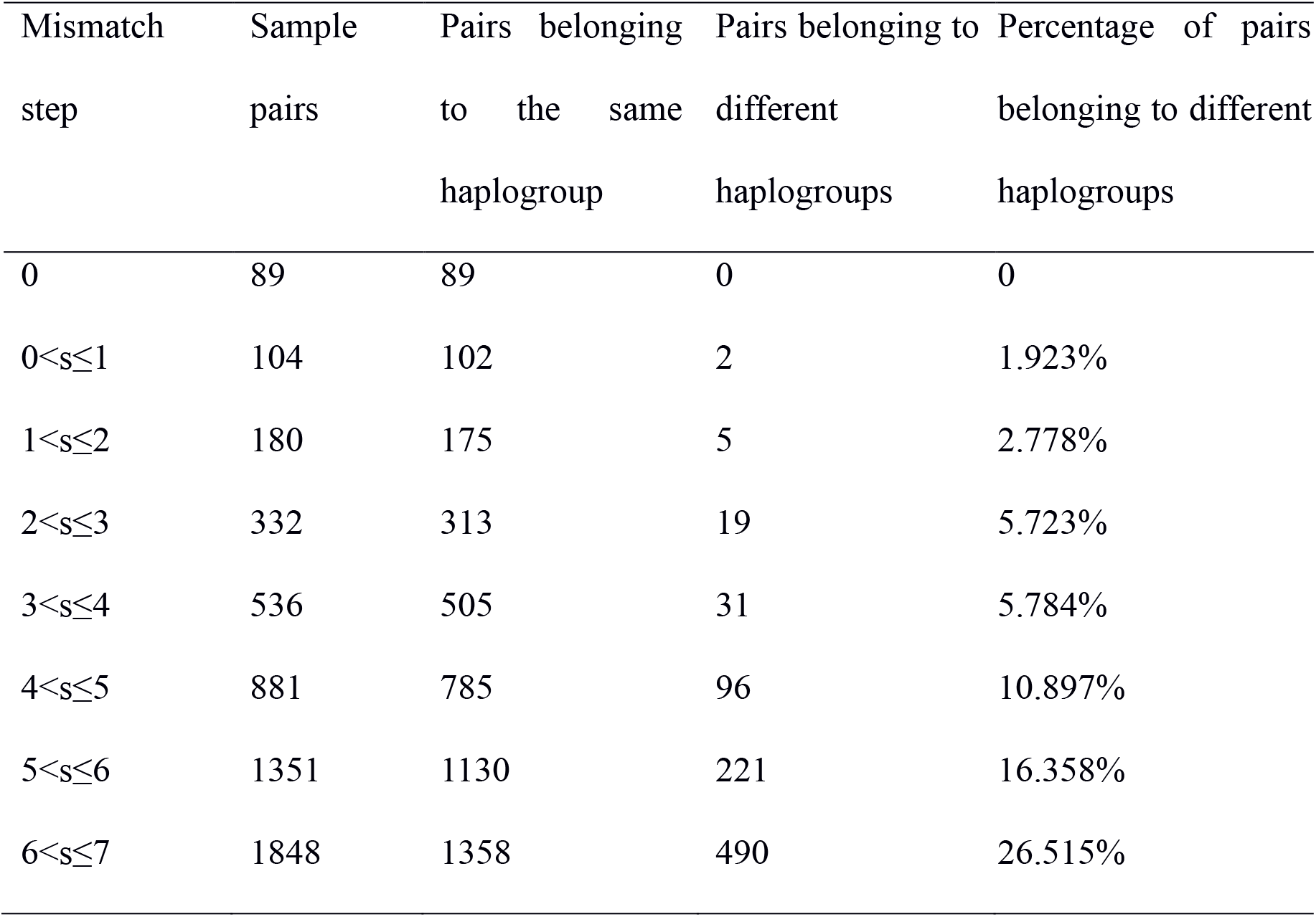
Indications from mismatch step results (mismatch step is the total number of different allele steps). Sample pairs are the number of pairs in the corresponding mismatch step.

| Mismatch step | Sample pairs | Pairs belonging to the same haplogroup | Pairs belonging to different haplogroups | Percentage of pairs belonging to different haplogroups |
| --- | --- | --- | --- | --- |
| 0 | 89 | 89 | 0 | 0 |
| 0<s≤1 | 104 | 102 | 2 | 1.923% |
| 1<s≤2 | 180 | 175 | 5 | 2.778% |
| 2<s≤3 | 332 | 313 | 19 | 5.723% |
| 3<s≤4 | 536 | 505 | 31 | 5.784% |
| 4<s≤5 | 881 | 785 | 96 | 10.897% |
| 5<s≤6 | 1351 | 1130 | 221 | 16.358% |
| 6<s≤7 | 1848 | 1358 | 490 | 26.515% |

